# Marker-Trait Complete Analysis

**DOI:** 10.1101/836494

**Authors:** Yi-Hui Zhou, Paul Gallins, Fred Wright

## Abstract

A recurring problem in genomics involves testing association of one or more traits of interest to multiple genomic features. Feature-trait squared correlations *r*^2^ are commonly-used statistics, sensitive to trend associations. It is often of interest to perform testing across collections {*r*^2^} over markers and/or traits using both maxima and sums. However, both trait-trait correlations and marker-marker correlations may be strong and must be considered. The primary tools for multiple testing suffer from various shortcomings, including *p*-value inaccuracies due to asymptotic methods that may not be applicable. Moreover, there is a lack of general tools for fast screening and follow-up of regions of interest.To address these difficulties, we propose the MTCA approach, for <u>M</u>arker-<u>T</u>rait <u>C</u>omplete <u>A</u>nalysis. MTCA encompasses a large number of existing approaches, and provides accurate *p*-values over markers and traits for maxima and sums of *r*^2^ statistics. MTCA uses the conditional inference implicit in permutation as a motivational framework, but provides an option for fast screening with two novel tools: (i) a multivariate-normal approximation for the max statistic, and (ii) the concept of eigenvalue-conditional moments for the sum statistic. We provide examples for gene-based association testing of a continuous phenotype and cis-eQTL analysis, but MTCA can be applied in a much wider variety of settings and platforms.

## 2 Introduction

Issues in multiple testing arises in a number of settings, with considerations of power and maintaining appropriate error control. Problems arising in genetic association especially benefit from careful consideration, as correlation structures may be strong, so that simple methods for family-wise error control may be overly conservative, and statistics that aggregate signal over multiple features may plausibly improve power.

These considerations have given rise to a number of purpose-built methods and associated software for genetic analysis. A non-exhaustive list includes methods for gene-based association testing of multiple markers with a disease trait [8], rare-variant association statistics in which multiple markers are aggregated [10], region-based eQTL association analyses [5], pathway associations of genetic markers or expression vs. a trait, analysis of associations with multiple transcriptional isoforms [9], and association analysis of genetic markers with multiple possibly correlated traits.

For the array of data types listed, often the solutions have been specific and purpose-built. For example, methods such as GATES [8] have been developed for gene-based association testing, [7] for multiple-trait association, [11] for gene-based cis-eQTL analysis, sQTLseeker [9] for isoform eQTL analysis, etc. Many existing software packages accordingly have specific input requirements. There are often good reasons to seek problem-specific solutions, for example if likelihood methods can offer improvements in power over trend testing. In addition, reliance on parametric assumptions may be convenient and appropriate in some settings, while resampling methods might be necessary in other settings.

In this paper we emphasize the commonalities in the analysis problems described, rather than the differences, and argue that a common data structure applies for many disparate problems. We provide software for analysis as an R package, and the results can be easily compared with the results of purpose-built software. Indeed, our work is partly motivated by our own desire to easily follow up on results from existing methods, for example in determining which feature among many is responsible for the signal in a significant result, or whether a significant finding is highly reliant on parametric assumptions or would also be supported by resampling approaches.

Here we present <u>M</u>arker-<u>T</u>rait <u>C</u>omplete <u>A</u>nalysis (MTCA), an analysis platform for multiple trend testing for a variety of genetic analyses. The approach provides family-wise false positive control over features tested (i.e. the maximum statistic), and also provides false-positive control for aggregated (summed) measures of association. Permutation methods are available as a default basis for testing, and are further enhanced with density fitting to enable estimation of highly significant *p*-values. Finally, we have developed novel analytic methods to provide fast approximate error control, in settings where permutation would be time-consuming.

### 2.1 A common analysis setup

We define 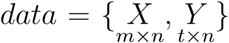 as two matrices to be compared for *m* genetic markers and *t* traits for *n* individuals. Typically *data* corresponds to a single genomic region (for the markers), with *mt* < 10, 000. The terms “marker” and “trait” are used for convenience, but the approach is entirely generic. We are interested in trend associations between *X* and *Y*, measured as the *m* × *t* matrix *r*^2^ of squared correlations between rows of *X* vs. rows of *Y*. Our null hypothesis is that *X* and *Y* were generated independently. We list a few important examples:

- Genetic association analysis of one or more correlated genetic markers (often representing a gene or pathway) versus *t* correlated disease traits.
- Regression-based cis-eQTL analysis comparing multiple SNP markers (*m* typically 1000 or fewer) to a single gene’s expression trait (*t* = 1).
- cis-eQTL analysis of multiple transcript isoforms (available from RNA-Seq data) *m* nearby markers to *t* individual exon counts (*t* typically in the range 3-20) for a gene.

For many problems of interest, our analysis will be performed separately for each *data*_*g*_ for *g* = 1, …, *G*, where typically *g* corresponds to a region on the genome, and *m* and *t* may also vary with *g*. Results for each *data*_*g*_ will be passed for downstream analysis using standard methods such as false-discovery control, and so for most of our development we focus our attention on a single *data*_*g*_ and suppress the *g* subscript.

Some of the motivation and individual components within MTCA are anticipated in other work, and in particular direct permutation methods are widely performed for some of the statistics proposed, such as gene-based cis-eQTL detection [5]. However, direct permutation is often computationally intensive to achieve sufficient resolution for relative ranking of highly significant results. Moreover, some existing methods perform family-wise error control over multiple features but do not aggregate, while others may only aggregate. We view our approach as providing a general purpose, convenient, and accurate screen for associations, and after screening across multiple {*data*_*g*_}, our software enables quick follow-up exploration of the source of signal. In addition, our development includes novel derivations for extremely rapid screening that is useful for large *G*. A full MTCA analysis provides a comprehensive output that is useful as a one-stop summary of association evidence.

## 3 Data structure and an illustration

### 3.1 The MTCA approach

We will consider control of family-wise error using maxima of statistics over markers and traits, as well as aggregate statistics that sum across rows and columns of *r*^2^, accounting for correlation structures within each of *X* and *Y*. In addition, we will consider grand summaries over the entire *data*. The computational effort for MTCA grows with both *m* and *t*, but again this is employed for a single region at a time and the computation is manageable. We assume that trend associations between *X* and *Y* are of interest. For a large variety of problems, the Pearson correlation *r* is powerful as a measure of association, either based on linear model assumptions or as efficient score statistics for nonlinear or generalized linear models. An exhaustive consideration of covariate control is beyond our scope here. However, we note that if normal linear modeling of the form *Y* ∼ *X* + *Z* is envisioned for covariate *Z*, then we can first linearly pre-residualize both *X* and *Y* for covariate *Z*. The correlations between *X*_*resid*_ and *Y*_*resid*_ are then partial correlations from the full linear model, and we simply use these residual matrices as our data. Figure 1 depicts both the data and the summarized evidence in a schematic.

**Figure 1:**
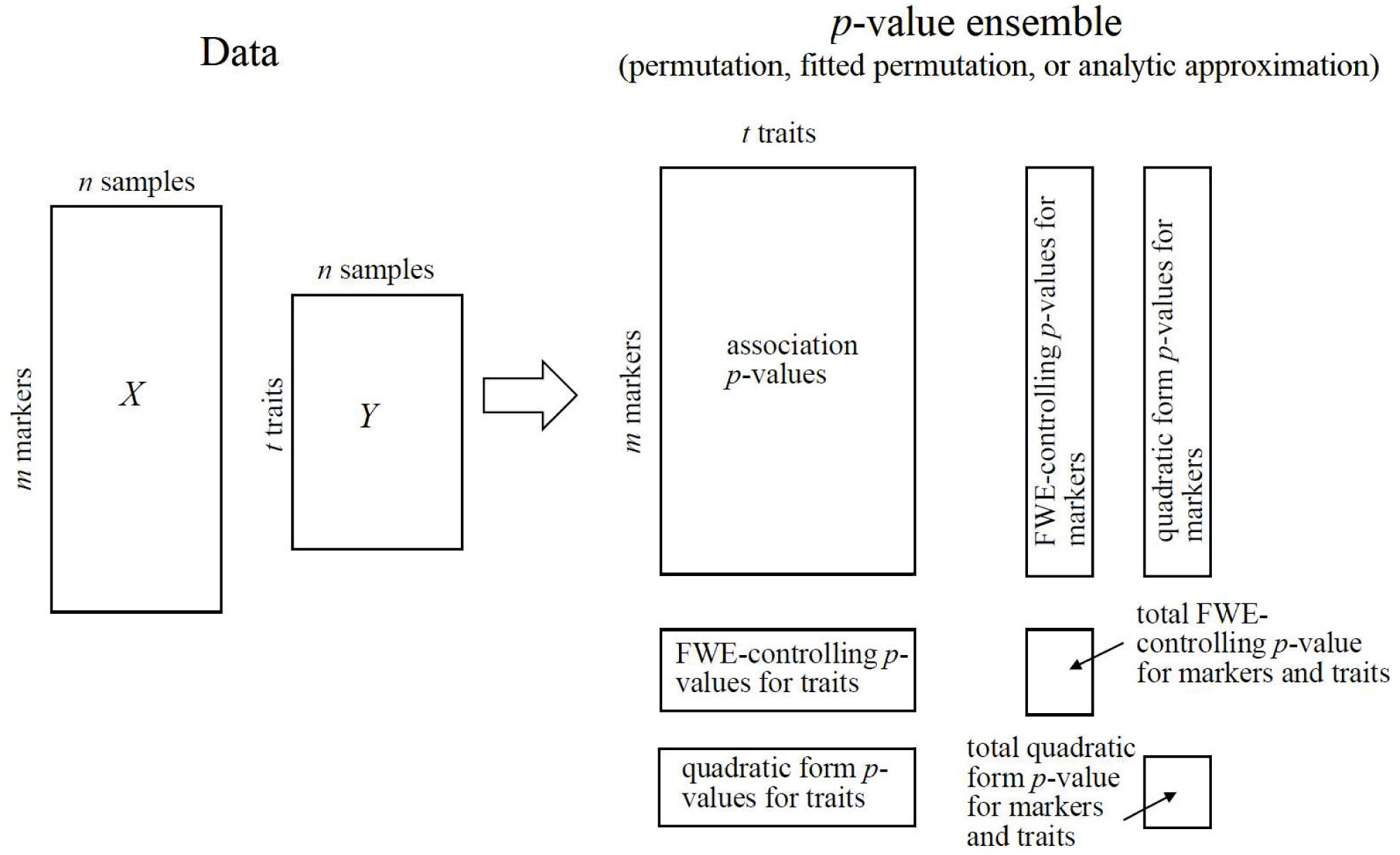
A schematic view of Marker-Trait Complete Analysis. The generic data structure is commonly encountered, and the *m* × *t* association *p*-values are straight-forward to compute. However, the maxima and sums are affected by correlation structures among rows of *X* and of *Y*, and *p*-values in grey are computed using permutation or novel approximation methods introduced in this paper.

The association evidence is summarized in the *m* × *t* matrix of *p*-values for squared correlations 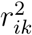 for each marker *i* and trait *k*. We define the max statistics over markers and traits as 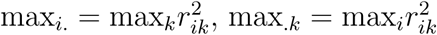. The analogous sum statistics are 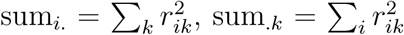. Braces {} are used to indicate a collection of indices, so e.g. sum_{*i*}_ is the vector of sum statistics over the markers. Finally, the total statistics over markers and traits are 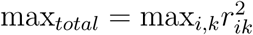 and 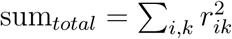.

The max_{*i*}_ vector represents evidence that the markers affect at least one trait, while sum_{*i*}_ is an aggregate statistic sensitive to association of multiple traits to the marker. The column-wise statistics max_{*k*}_ and sum_{*k*}_ provide analogous information across markers. Both the maxima and sums are of interest in a variety of genetics problems. The total statistics have a pleasing symmetry, where max_*total*_ = max(max_{*i*}_) = max(max_{*k*}_) and sum_*total*_ = sum(sum_{*i*}_) = sum(sum_{*k*}_).

Computing accurate *p*-values for the statistics requires consideration of correlations among the markers and traits, and below we describe methods to obtain these *p*-values. The column and row *p*-values offer an immediate summary of the degree of evidence for association within the respective marker or trait, after accounting for correlation structures. We also compute total *p*-values for the grand maxima and sum of the *r*^2^ matrix. Collectively, we refer to the *p*-value objects as the “*p*-value ensemble.”

*P*-values from the maximum statistic control family-wise error for the collection being tested. As we show later, the sum statistic for rows and columns is a quadratic form. In settings where numerous {*data*_*g*_} are screened, we expect *p*_*g,max total*_ or *p*_*g,sum total*_ to be the primary basis for screening. Any significant findings, after FDR control over *G* tests, can be followed up with post-hoc analysis of the *p*-value ensemble for significant *data*_*g*_.

MTCA can compute three types of *p*-value ensembles (Figure 1). The first (permutation) consists of empirical *p*-values based on direct permutation of columns of *Y* compared to *X*. From the permutations, density fitting to the max *r*^2^ and sum *r*^2^ (permutation + fitting) can be performed for little additional computational cost, providing enhanced resolution for extremely small *p*-values. However, the results may be viewed as somewhat dependent on distributional assumptions. A third *p*-value ensemble (analytic approximation) consists of approximate *p*-values based on novel methods described later. These approximations may be attractive when the investigator wishes to keep the computational cost low for each {*data*_*g*_}, for example when screening a genome.

### 3.2 An illustration

Before describing the methods to obtain *p*-value ensembles, we illustrate with a simulated example to guide the reader in interpretation. We simulated a dataset with *m* = 200 markers (*X*) and *t* = 10 traits (*Y*) for *n* = 100 individuals. The markers were simulated to be similar to SNP genotypes (values 0, 1, or 2) with minor allelele frequency 0.2, in 10 blocks of varying size and high correlations (in excess of 0.9) between successive markers. Trait 1 was simulated to depend on marker 1 as *y*_1_ = *βx*_1_ + ∊, where ∊ ∼ *N* (0, 1 − *β*^2^) for *β* = 1/2. Traits 3-8 were simulated in the same manner as all depending on marker 100, but with *β* = 1/6. Finally, trait 10 was simulated as depending on marker 150 with *β* = 1/2.

Figure 2 shows an MTCA plot of the permutation+density *p*-value ensemble based on 10,000 permutations. *P*-values for the *r*^2^ matrix are shown in heatmap form, to see the marker-trait pairs with greatest association. For both rows and columns, “mini-Manhattan” plots show the corresponding max *r*^2^ and sum *r*^2^ *p*-values. To aid interpretation, the points are colored red if they have an FDR *q* < 0.05 after correction for the points in each mini-Manhattan plot. Finally, the *p*_*max total*_ and *p*_*sum total*_ are provided, along with 95% confidence intervals for these quantities.

**Figure 2:**
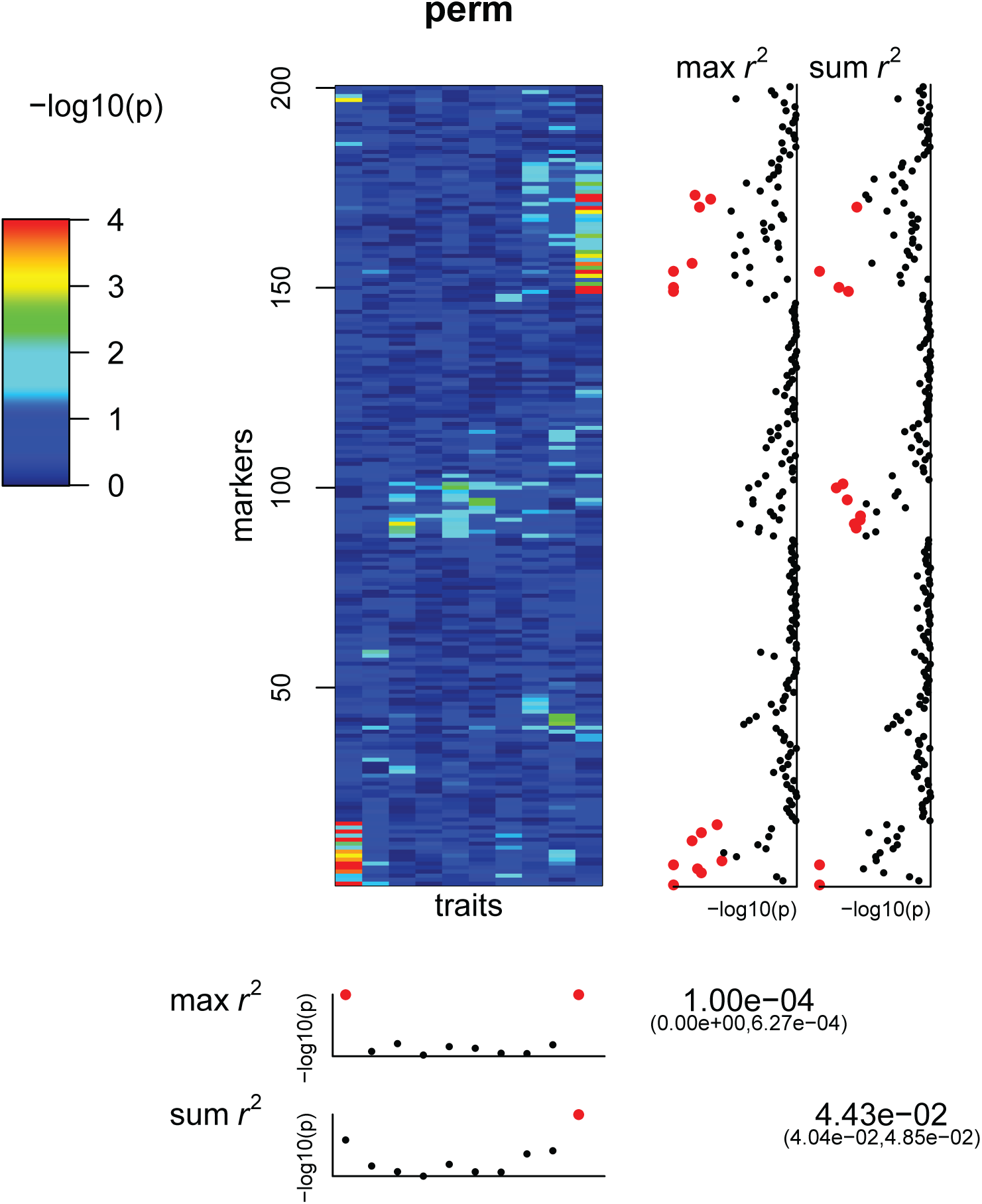
Example MTCA plot for a *p*-value ensemble using 10,000 permutations, for 200 markers and 10 traits, with simulated true effects at markers 1 (first trait), marker 100 (traits 3-8) and marker 150 (10th trait). *P*-values shown in red have FDR *q*-value < 0.05 for the corresponding mini-Manhattan plot.

The sources of signal in the data are apparent. The first several markers show evidence of association with trait 1 in both types of marker mini-Manhattan plots, as do the markers near marker 100 for trait 10. Groups of nearby markers have *q* < 0.05 due to marker correlation structure. As only a single trait is associated with each respective region, it is reasonable that more findings are made for the max statistic than the sum statistic. For the weaker signal at marker 100, only the sum statistic identifies associations for the marker mini-Manhattan plot, as it is able to aggregate evidence across several traits. The trait max *r*^2^ mini-Manhattan plot shows associations for both traits 1 and 10. Finally, the total *p*-value for max *r*^2^ provides evidence for overall associations in the full matrix, while the total *p*-value for max *r*^2^ is barely significant at *α* = 0.05.

## 4 Methods

### 4.1 Permutation

Under the null hypothesis, *X* is independent of *Y*, and the *n*! arrangements of columns of *Y* relative to *X* are equally likely. MTCA computes *r*(*X, Y*) as the *m* × *t* Pearson correlation matrix upon which all summaries are based. Let Π denote a random permutation of the indices {1, …, *n*}, chosen uniformly from the *n*! possibilities, so that *r*(*X, Y*_Π_) is the random correlation matrix where columns of *Y* are re-ordered according to Π. Under the null hypothesis, permutation provides a valid null population, without requiring distributional assumptions or a generating model for *X* and *Y*. We use *s*(*X, Y*) to generically represent our statistic (maximum or sum, over columns, rows, or total matrix), and suppose *H* random permutations are performed, indexed *h* = 1, …, *H*. The permutation *p*-value estimate is then 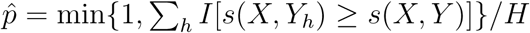, computed for all of the row, column and total statistics. These permutation *p*-values are obtained for all seven data objects in the *p*-value ensemble (Figure 1). MTCA also reports Agresti-Coull confidence intervals [1] for the permutation *p*_*total*_ values, so that users may judge whether the accuracy is sufficient for their purpose.

Setting the minimum *p*-value to 1*/H* is necessary to avoid exaggeration of significant findings, and the upper confidence limit will always exceed 1*/H*. For a single *data*_*g*_, it may be feasible and desirable to perform a large number (say 10^5^) permutations to obtain sufficient resolution for downstream inference. However, when *G* is also large, extensive permutation presents difficulties in computation. Moreover, it is often of interest biologically to distinguish among the most highly significant findings, but difficult if they all achieve the same minimum value of 1*/H*.

#### Adaptive permutation

One straightforward way to minimize computation is to perform adaptive permutation, in which few permutations are needed if it is clear that *p*-values will not be significant. Numerous existing methods have been developed for sequential *p*-value estimation. However, most methods (e.g. [3]) are intended to avoid errors in significance testing relative to a fixed threshold *α*, while the downstream uses of the MTCA *p*-value ensembles will vary, depending on the problem. We have thus we have implemented a simple criterion, based on relative accuracy of the total *p*-value estimates to their binomial standard errors. Starting with a specified minimum *H*_*min*_, permutation is halted if 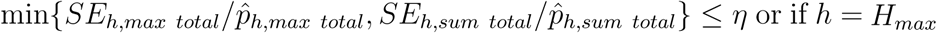. A typical implementation in our examples uses *H*_*min*_ = 1000, *η* = 0.1, and *H*_*max*_ = 10, 000. During the permutation resampling, MTCA saves all of the permuted row and column max *r*^2^ and sum *r*^2^ values, which are necessary for the density fitting below. We note that there is no need to save all the permutations of the larger *r*^2^ matrix itself − MTCA computes permutation *p*-values for the individual marker-trait associations merely by keeping a running total of the number of permuted associations that exceed each data element in the original *r*^2^ matrix.

### 4.2 Permutation-based density fitting

To address the difficulty in ranking *p*-values for highly significant associations, we implement density fitting to the max *r*^2^ and sum *r*^2^ statistics. The basic idea is to use the permutations to determine empirical moments, followed by fitting of an appropriate density to the distributions. The steps below assuming we are computing a row (marker) statistic, but are the same whether we are dealing with a row statistic, a column statistic, or a total statistic. For the observed data and for each permutation *h* for max *r*^2^, we first compute standard correlation *p*-values, *p*_*obs*_ and *p*_*h*_, using the assumed null distribution 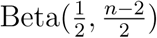. The resulting values over permutations tend to be left-shifted compared to uniform, because we have taken maxima over (usually multiple) features. From these *H p*-values we then use the empirical mean and variance over permutation to estimate the parameters of a corresponding beta distribution function 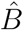, reflecting the left-shift in the values. Finally, a corrected *p*-value is 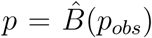. For sum *r*^2^, we instead work directly with the sums and use a 4-parameter Pearson family density approximation using the `PearsonDS` package [2] to the first four empirical moments of the permutation distribution to obtain a *p*-value.

MTCA reports *p*-values based on density fitting for all the objects in the *p*-value ensemble. For the individual elements of the *r*^2^ matrix, null beta assumptions are used and don’t require saving permutations for density fitting.

### 4.3 Analytic approximations

Analytic approximate *p*-values here are motivated by using the permutation null distribution, which conditions on the observed *X* and *Y*. The use of the permutation null helps clarify certain aspects of the analysis, such as the relevance of sample data moments, which often determine the moments of statistics under the *population* of all possible permutations. In the discussion below, for definiteness, we describe results for the maximum and sum statistics over markers, and the same approach may be used over traits.

#### The max statistic

To compute *p*-values for the maximum statistic, we first establish an under-appreciated fact about correlation coefficients under permutation. Using the notation above, the random (under permutation) Pearson correlation coefficients for features *x*_1._ and *x*_2._ are *r*(*x*_1._, *y*_Π_) and *r*(*x*_2._, *y*_Π_), and the correlation of correlations is *corr*(*r*(*x*_1._, *y*_Π_), *r*(*x*_2._, *y*_Π_)) = *r*(*x*_1._, *x*_2._) (Appendix **1.2** of Supplementary file). In other words, the true permutation correlation matrix of the *r* vector is exactly the observed sample correlation matrix over markers. This correlation result, in combination with a normal approximation to the Fisher-transformed *r*, is used to provide the mean and variance for the number of instances of *p*-values < *α* for fixed *α* (Appendix **1.3** of Supplementary file). Finally, an approximate family-wise error-controlling *p*-value is obtain by fitting a discrete distribution to this computed mean and variance. The same approach can be used for the max_*total*_ statistic (Appendix **1.4** of Supplementary file) using a Kronecker product of correlation matrices for *X* and *Y*, made faster using a correlation-binning approach and a fast approximation to the bivariate normal cdf.

#### The sum statistic

For the sum statistic, we use the general strategy of moment-fitting approaches applied in [14] and [12], followed by Pearson family density fitting to obtain *p*-values. For high accuracy, the first four exact permutation moments can be computed using quadratic form results from [14], including a novel formulation detailed in Appendix **1.7** of Supplementary file, as sum_*total*_ is not a quadratic form. However, this general strategy becomes less attractive for large *n*, as the computation of these exact moments is *O*(*n*^3^).

To develop a faster approach, we note that the beta distribution approach of [12] enabled approximation of the first two moments of the sum statistic using only the eigenvalues of *X*. Briefly, from the fact that Σ_*i*_ *λ*_*i*_ = *m* we have 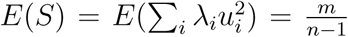, and by using covariance results from [12] we obtain 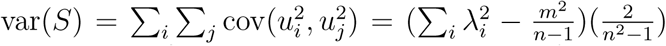.In Appendix **1.8** of Supplementary file, we derive the third and fourth moments, with 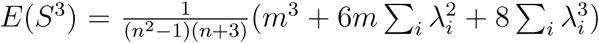 The fourth moment requires more effort, and can be shown after several cancellations to be 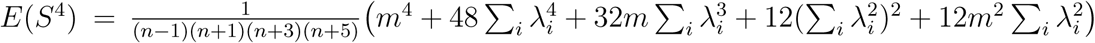. Collectively, the four moments enable accurate Pearson family density fitting for row and column sum statistics, although the grand sum_*total*_ requires an additional approximation. We term the approach as *eigenvalue* conditional, and can be viewed as motivated by sampling from multivariate normal *X* or *Y*, but by conditioning on the observed sample eigenvalues we consider the approach as applicable to the permutation null. We note that the approach is entirely scalable to various combinations of *m* and *n*, and in particular has no requirement that *m* < *n*. In this manner our approximation has distinct advantages over quadratic form approximations that require large *n*.

### 4.4 Simulation conditions and real datasets

The maximum and sum statistics are familiar and used in numerous contexts. Thus we are not concerned here with the *power* of the statistics, which are already in common use, but in demonstrating proper false positive control. We performed simulation of datasets under a range of conditions, and show the results from analysis of two real datasets.

#### Simulation

We simulated 100,000 instances of all 1296 combinations of the following conditions:

- *n* = {50, 500}
- *m* = {10, 100, 1000}
- *t* = {2, 6, 10}
- *X* was either multivariate normal *MV N* (0, Σ_*X*_), with covariance matrix Σ described below, or discrete (taking the values {0, 1, 2}) to mimic SNP genotype data. The discrete data were produced by starting with multivariate normal data *X*′ ∼ *MV N* (0, Σ_*X*_) and thresholding *X* = *f* (*X*′). Here the function *f* followed the procedure of [13], thresholding the normal data to produce genotype-like count data that accords with minor allele frequencies (MAFs) observed in GWAS data, with average MAF of about 0.2.
- *Y* was either multivariate normal *MV N* (0, Σ_*Y*_), with covariance structure described below, or binary by starting with *Y*′ ∼ *MV N* (0, Σ_*Y*_) data, and thresholding *Y* = *I*(*Y*′ > 0)
- The covariance matrix Σ_*X*_ has diagonal 1, and is chosen to be either (i) block diagonal with 10 equal-sized blocks, with within-block covariance (correlation) *ρ*_*X*_; (ii) consisting of a large block of *m/*2 markers with covariance *ρ*_*X*_ and the remaining markers uncorrelated.
- The covariance Σ_*Y*_ similarly depends on *ρ*_*Y*_ for *t* traits. Due to the smaller range of *t*, only the “large block” option is used.
- *ρ*_*X*_ = {0, .5, 0.8} Note that only positive correlations need be considered, because we are working with squared correlations *r*^2^. Also, when *ρ* = 0, there is no block structure to the covariance matrix.
- *ρ*_*Y*_ = {0, .5, 0.8}

As *X* and *Y* are independent, permutation is guaranteed to provide rank-uniform *p*-values. We focus on the total statistics, as these are the most difficult to approximate, and record the rejection probabilities for fitted permutation and analytic approximations under target *α* = 0.05.

#### Real datasets

The first dataset represents 5663 patients from the Cystic Fibrosis (CF) International Gene Modifier consortium, in which genotypes at 8 million markers (mostly imputed) were compared to phenotypes. Among the most important associations are with two quantitative traits: a measure of lung function [6] and martingale residuals from time-to-onset of CF-related diabetes [4]. Both markers and traits were pre-residualized for the covariates (including cohort sex and genotype principal components) used in the original analysis. Here we focus on a single candidate gene EEA1 (1093 markers within 100 kb of transcription start site) merely to illustrate the approach and the behavior of our statistics.

The second dataset consists of 97 individuals contributing liver eQTL data for the GTEx Consortium [5], with RNA-Seq read counts compared to SNP genotypes. Here cis-effects for each gene *g* were assessed by constructing *data*_*g*_ using all SNPs within a 100 kb window of the transcription start site. Analyses were performed using overall expression for each gene as a quantitative phenotype, as well as exon-specific analysis. The exon-specific analysis first divided the read counts for each exon by the overall read count (gene-wide), thus measuring how exon “usage” is affected by genotype. These exon-specific measurements offer an excellent example of a collection of “traits” with inherent dependence (they must sum to 1 after our scaling) can be examined collectively. We focus on the gene RPS28, which was among the most significant in a genome-wide scan for eQTL effects.

## 5 Results

Table 1 shows the Type I error results for the analytic approximation for the max_*total*_ statistic, over a representative number of the conditions. The family-wise error control is near nominal, except when *ρ*_*Y*_ = 0 and *ρ*_*X*_ = 0.8, where the results are somewhat conservative. Table 2 shows Type I error results for the sum_*total*_ statistic, and is slightly anticonservative for large *ρ*_*X*_. The full results for the 1296 conditions are available as Supplementary files. In addition, the Supplementary files include the results for the permutation+fitting approach, which shows near nominal Type I error control throughout.

**Table 1:**
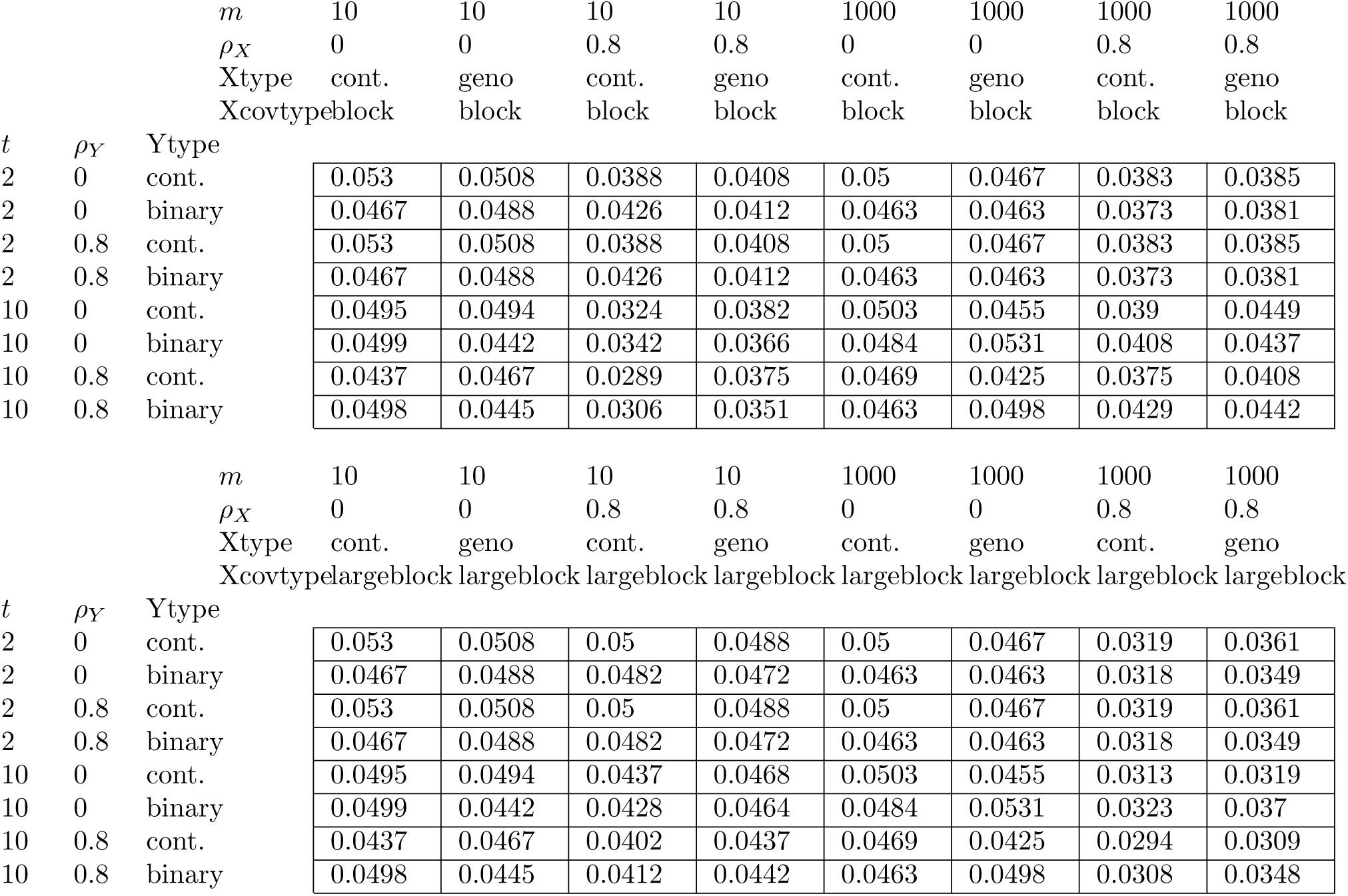
Analytic max *r*^2^, *n* = 50.

**Table 2:**
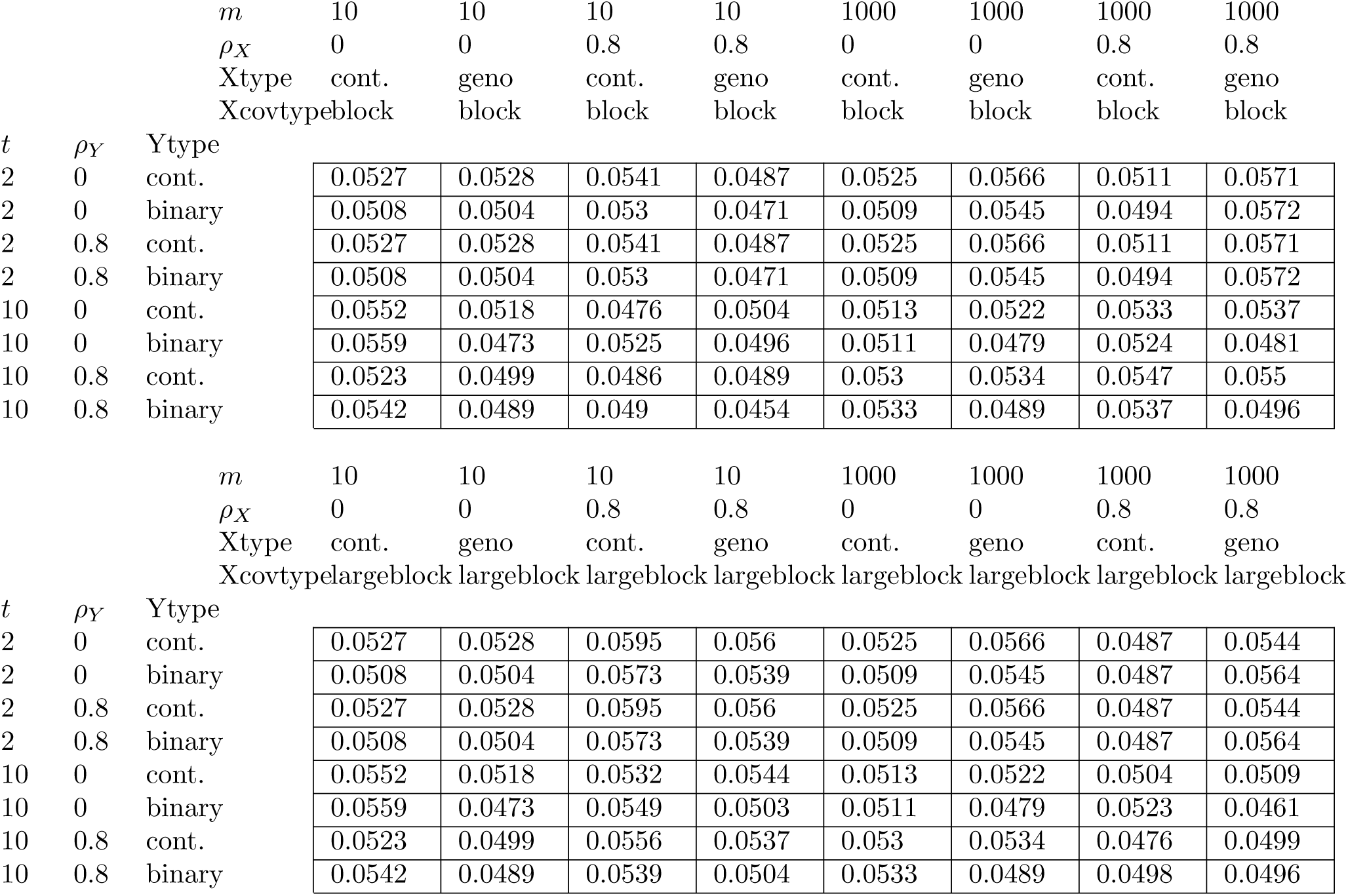
Analytic sum *r*^2^, n = 50.

Figure 3 shows the CF results for markers near EEA1 for the two phenotypes. The *p*-value ensemble (panel A) is nominally significant at *α* = 0.05 for the first phenotype (lung function), but not the second (the diabetes phenotype), both in terms of FDR-controlling values and for the total statistics. Although the results do not meet multiple testing standards, we are primarily interested in using the data to illustrate type I error control. Panel B shows the results for 10,000 permutations of the data, with the sum statistic for rows, columns, and total shown. Each row and column provides a separate realization, so there are for example *m* × 1, 0000 elements in the row histogram. Overlaid on the histograms is the Pearson family density fit, and *qq*-plots of the −log_10_ *p*-values are shown, all of which show good type I error control.

**Figure 3:**
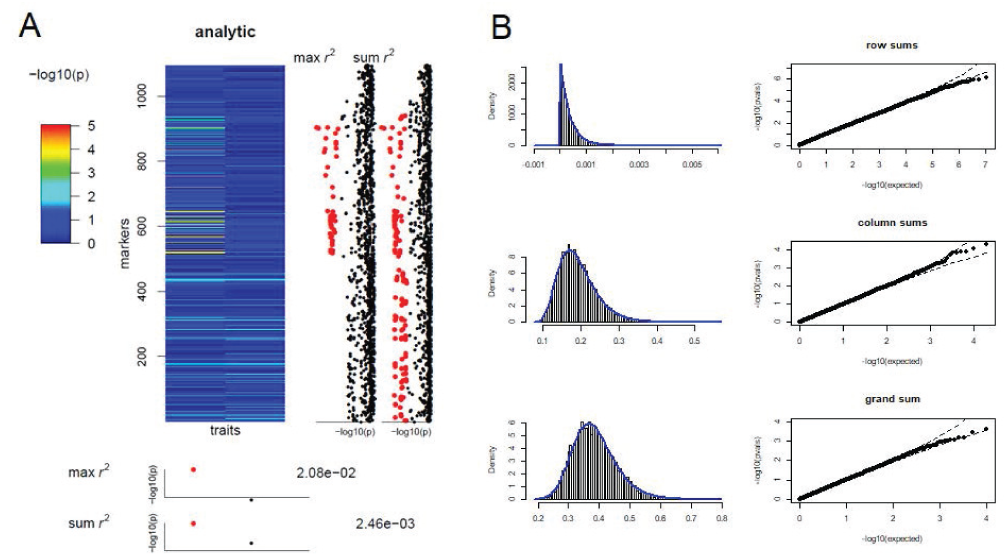
EEA1 example for cystic fibrosis lung and diabetes traits. (A) *p*-value ensemble (B) For the sum *r*^2^ statistics, histogram of permutations overlaid with analytic density fits and corresponding *qq* plots.

Figure 4 shows results for gene RPS28 for the liver GTEx eQTL data. Panel A shows a Manhattan plot for fast analytic approximate *p*-values of the Σ_*total*_ statistic. Gene RPS28 is indicated, and is among the most significant genome-wide. The *p*-value ensemble for total gene expression (panel B) shows very high significance for both the max and sum statistics, which would survive even the most stringent error control procedure across genes. Panel C shows the *p*-value ensemble for exon usage (only three exons had sufficient read counts). The evidence for differential exon usage is much less significant, but still striking (*p* = 3.04 × 10^−5^ for the sum statistic). Panel D shows the extremely strong relationship of total gene expression to allele counts for marker rs2972574, while panel E shows the average relative exon usage across the exons for the three genotype values. The results show a very low relative exon usage for exon 1 throughout, and increasing number of minor alleles shows a modest relative increase in usage of exon 3 at the expense of exon 2.

**Figure 4:**
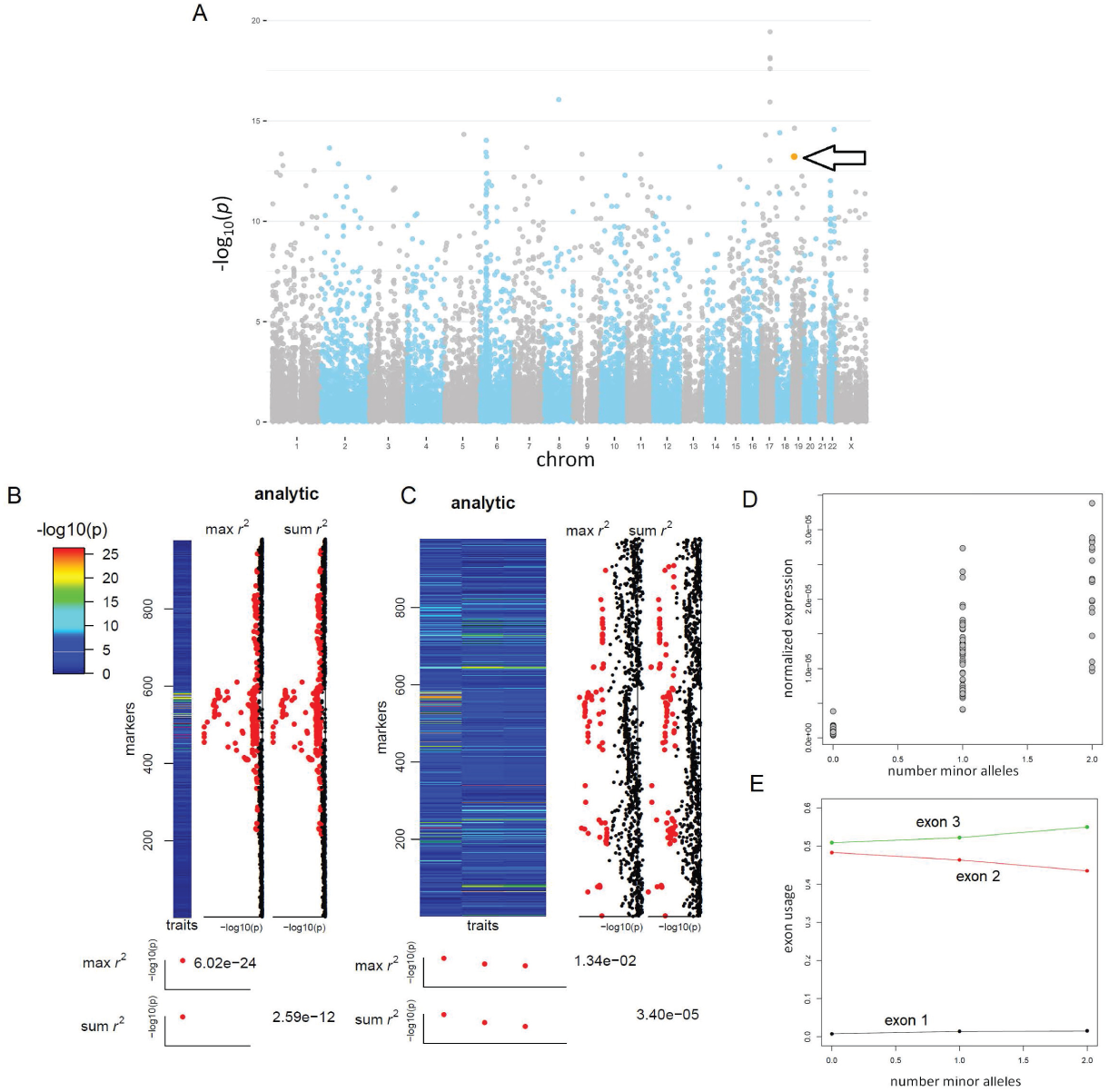
Liver eQTL plot. (A) Manhattan plot for analytic sum_*total*_, with RPS28 highlighted. (B) *p*-value ensemble for total gene expression. (C) *p*-value ensemble for exon usage, with relative expression of exons 1-3 as “traits.” (D) Overall expression vs. number of minor alleles of SNP rs2972574. (E) For the same marker, relative exon usage for the three exons.

**Figure 5:**
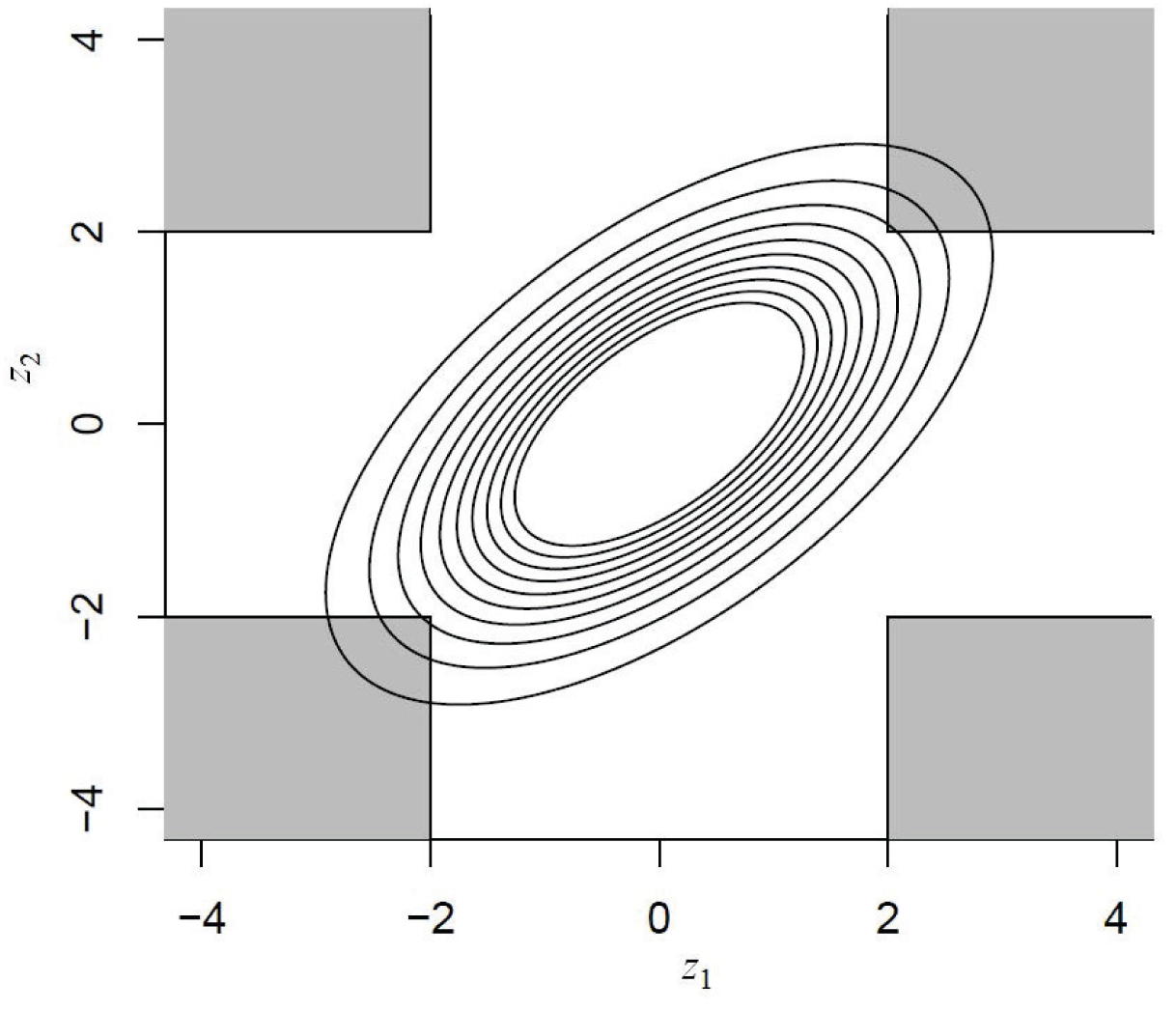
For the standard bivariate normal density with *ρ* = 0.6, the shaded portion identifies the region for which *I*_1_ = 1, *I*_2_ = 1 for threshold *z*_*α*_ = 2.

## 6 Discussion

We have presented MTCA, a comprehensive method for screening a genome for associations of sets of markers with one or more traits, accounting for correlation structures inherent in the data. The use of *p*-value ensembles is potentially very valuable, enabling researchers to display and further “drill down” into the data in a systematic fashion. In addition, the use of both max and sum statistics ensures sensitivity to a range of alternatives and types of genomic signals.

We propose MTCA as a general-purpose tool, and the availability of an *R* package will make it useful in a number of association studies. As a general tool, MTCA is not intended to be annotation-aware, or to replace any number of purpose-built bioinformatics tools. For sequence-based association, we expect that the analytic approximation may be less accurate, and the user may wish to use permutation+fitting to obtain *p*-values.

## 7 Supplementary Material

### Funding

Funded in part by an internal GRIP award at NC State University, a CF Foundation grant, and NIH grants R01HG009125 and R01ES029911. For the CF data, we gratefully acknowledge the CF patients, the Cystic Fibrosis Foundation, the UNC Genetic Modifier Study, the Canadian Consortium for Cystic Fibrosis Genetic Studies, funded in part by Cystic Fibrosis Canada and by Genome Canada through the Ontario Genomics Institute per research agreement 2004-OGI-3-05, with the Ontario Research Fund—Research Excellence Program, and the French CF Gene Modifier Study, funded in part by the French CF foundation Vaincre La Mucoviscidose.

## References

[1] A. Agresti and B. A. Coull. Approximate is better than “exact” for interval estimation of binomial proportions. The American Statistician, 52(2):119–126, 1998.

[2] M. Becker, S. Klößner, and M. M. Becker. Package ‘pearsonds’. Aust. NZJ Stat, 50(2):199–205, 2017.

[3] J. Besag and P. Clifford. Sequential monte carlo p-values. Biometrika, 78(2):301–304, 1991.

[4] S. M. Blackman, C. W. Commander, C. Watson, K. M. Arcara, L. J. Strug, J. R. Stonebraker, F. A. Wright, J. M. Rommens, L. Sun, R. G. Pace, et al. Genetic modifiers of cystic fibrosis–related diabetes. Diabetes, 62(10):3627–3635, 2013.

[5] G. Consortium et al. Genetic effects on gene expression across human tissues. Nature, 550(7675):204, 2017.

[6] H. Corvol, S. M. Blackman, P.-Y. Boëlle, P. J. Gallins, R. G. Pace, J. R. Stonebraker, F. J. Accurso, A. Clement, J. M. Collaco, H. Dang, et al. Genome-wide association meta-analysis identifies five modifier loci of lung disease severity in cystic fibrosis. Nature communications, 6:8382, 2015.

[7] J. Kim, Y. Zhang, and W. Pan. Powerful and adaptive testing for multi-trait and multi-snp associations with gwas and sequencing data. Genetics, 203(2):715–731, 2016.

[8] M.-X. Li, H.-S. Gui, J. S. Kwan, and P. C. Sham. Gates: a rapid and powerful gene-based association test using extended simes procedure. The American Journal of Human Genetics, 88(3):283–293, 2011.

[9] J. Monlong, M. Calvo, P. G. Ferreira, and R. Guigó. Identification of genetic variants associated with alternative splicing using sqtlseeker. Nature communications, 5:4698, 2014.

[10] M. C. Wu, S. Lee, T. Cai, Y. Li, M. Boehnke, and X. Lin. Rare-variant association testing for sequencing data with the sequence kernel association test. The American Journal of Human Genetics, 89(1):82–93, 2011.

[11] H.-C. Yang, C.-W. Lin, C.-W. Chen, and J. J. Chen. Applying genome-wide gene-based expression quantitative trait locus mapping to study population ancestry and pharmacogenetics. BMC genomics, 15(1):319, 2014.

[12] Y.-H. Zhou, W. T. Barry, and F. A. Wright. Empirical pathway analysis, without permutation. Biostatistics, 14(3):573–585, 2013.

[13] Y.-H. Zhou, J. S. Marron, and F. A. Wright. Computation of ancestry scores with mixed families and unrelated individuals. Biometrics, 74(1):155–164, 2018.

[14] Y.-H. Zhou, G. Mayhew, Z. Sun, X. Xu, F. Zou, and F. A. Wright. Space–time clustering and the permutation moments of quadratic forms. Stat, 2(1):292–302, 2013.

