## Supplementary Materials for "Marker-Trait Complete Analysis"

BIOMETRICS 64, ??–?? DOI: 10.1111/j.1541-0420.2005.00454.x
December 2019

### Supplementary document for "Marker-Trait Complete Analysis"

**Yi-Hui Zhou**

Department of Biological Sciences, North Carolina State University, Raleigh, North Carolina, U.S.A

**and**

**Fred Wright**

Departments of Biological Sciences and Statistics, North Carolina State University,

Raleigh, North Carolina, U.S.A.

This paper has been submitted for consideration for publication in *Biometrics*

*Received October* 2019. *Revised February* 2019. *Accepted March* 2019.

#### Appendix

##### *Additional notation*

Define the sample mean of length-$n$ vector $x$ as $\bar{x} = \sum_j x_j/n$ and the sample standard deviation $s_x = \frac{1}{n-1}\sum_{j=1}^{n}(x_j - \bar{x})^2/(n-1)$. The Pearson sample correlation coefficient between $x$ and $y$ is $r(x,y) = \frac{\sum_j x_j y_j - n\bar{x}\bar{y}}{(n-1)s_x s_y}$. The population coefficient between random variables $X$ and $Y$ is $\mathrm{corr}(X,Y) = \frac{\mathrm{cov}(X,Y)}{\sigma_X \sigma_Y}$ where the denominator is the product of population standard deviations.

Our data consist of $m \times n$ marker data $X$ and $t \times n$ trait data $Y$, where column vectors are independently sampled. We use the complete null hypothesis that $X$ is generated independently of $Y$, although correlations among markers and among traits are arbitrary (and may be large). For illustration it is convenient to consider length-$n$ marker vectors $x_1, x_2$ (the first two rows of $X$) and length-$n$ phenotype vector $y$ (i.e. $t = 1$). Without loss of generality, we assume that each of the vectors has been pre-scaled so that, e.g., $\bar{x_1} = 0$, $s_{x_1} = \sqrt{1/(n-1)}$. Thus $r(x_1, x_2) = \sum_j x_{1j}x_{2j}$, $r(x_1, y) = \sum_j x_{1j}y_j$, etc.

For random permutation $\Pi$, let $\Pi[j]$ denote the $j$th index from permutation $\Pi$. A random Pearson correlation between $x_1$ and $y$ is $r(x_1, y_\Pi) = \sum_j x_{1j}y_{\Pi[j]}$. Although we consider each of $x_1$, $x_2$, and $y$ to be fixed, $r(x_1, y_\Pi)$ and $r(x_2, y_\Pi)$ are random variables constituting a population defined by the permutations. This permutation perspective is useful for motivation, and aligns our inference with that achieved by direct permutation. Another important consideration is that we permute entire columns of $Y$ (or equivalently, of $X$), so that both marker-marker and trait-trait correlations are preserved under permutation.

*The correlation between $r_\Pi$ at two markers equals the marker genotype correlation*

**Result 1**

$$\mathrm{corr}(r_{1,\Pi}, r_{2,\Pi}) = r(x_1, x_2), \tag{1}$$

i.e., the permutation *population* correlation between $r_{1,\Pi}$ and $r_{2,\Pi}$ is equal to the *sample* correlation between $x_1$ and $x_2$.

*Proof.* First, as shown in the safeExpress paper Zhou et al. (2013) it is helpful to note that $E(r(x, y_\Pi)) = 0$, $\mathrm{var}(r(x, y_\Pi)) = 1/(n-1)$ for any $x$ and $y$, provided they are not degenerate (i.e. both $x$ and $y$ must have positive sample variance). Also, we note that

$$\sum_j \sum_{j'} x_{1j} x_{2j'} = (\sum_j x_{1j})(\sum_{j'} x_{2j'}) = 0. \tag{2}$$

We have

$$\mathrm{cov}(\sum_j x_{1j} y_{\Pi[j]}, \sum_j x_{2j} y_{\Pi[j]}) = E(\sum_j x_{1j} y_{\Pi[j]} \sum_{j'} x_{2j'} y_{\Pi[j']}) = \sum_j \sum_{j'} x_{1j} x_{2j'} E(y_{\Pi[j]} y_{\Pi[j']})$$

$$= \sum_j x_{1j} x_{2j} E(y^2_{\Pi[j]}) + \sum_{j \neq j'} x_{1j} x_{2j'} E(y_{\Pi[j]} y_{\Pi[j']}). \tag{3}$$

Note that $E(y^2_{\Pi[j]}) = \sum_{j'} y^2_{j'}/n = s^2_y \frac{n-1}{n} = \frac{1}{n}$, because, for fixed $j$, the index $\Pi[j]$ is uniform on the integers $\{1, ..., n\}$. Also, for fixed $j \neq j'$, we have

$$E(y_{\Pi[j]} y_{\Pi[j']}) = E(E(y_{\Pi[j]} y_{\Pi[j']} | \Pi[j])) = E(y_{\Pi[j]} E(y_{\Pi[j']} | \Pi[j]))$$

$$= E(y_{\Pi[j]}(\frac{-y_{\Pi[j]}}{n-1})) = -E(y^2_{\Pi[j]})/(n-1) = -1/(n(n-1)).$$

The last three equalities follow from the fact that $\sum_j y_{\Pi[j]} = 0$ regardless of $\Pi$, so the average of $y_{\Pi[j']}$ over the $n-1$ instances of $j' \neq j$ must be $-y_{\Pi[j]}/(n-1)$. Plugging the results into (3), we have

$$\mathrm{cov}(r_{1,\Pi}, r_{2,\Pi}) = \sum_j x_{1j} x_{2j} (\frac{1}{n}) + \sum_{j \neq j'} x_{1j} x_{2j'} (\frac{-1}{n(n-1)})$$

$$= r(x_1, x_2)/n + (0 - \sum_j x_{1j} x_{2j})(\frac{-1}{n(n-1)}) = \frac{1}{n} r(x_1, x_2) + \frac{1}{n(n-1)} r(x_1, x_2) = \frac{r(x_1, x_2)}{n-1}, \tag{4}$$

and so finally $\mathrm{corr}(r_{1,\Pi}, r_{2,\Pi}) = \mathrm{cov}(r_{1,\Pi}, r_{2,\Pi})/(1/(n-1)) = r(x_1, x_2)$.

*Approximate FWER control over m markers using moments*

After Fisher transformation $z = \frac{1}{2}\log(\frac{1+r}{1-r})$ or other standardization, we consider each pair of statistics $\{z_{i\Pi}, z_{i'\Pi}\}$ as approximately bivariate normal under permutation, with correlation $r(x_i, x_{i'})$. Although result (1) technically applies to the correlations (not $z$), numerical investigations show the correspondence is very close, even for small $n$. For each marker we use the two-sided $p$-value $p_i = 2\Phi(-|z_i|)$, and define $I_i = I[p_i \leqslant \alpha]$ and $S = \sum_i I_i$. For nominal significance level $\alpha$, $P(\text{reject any hypothesis}) = 1 - P(S_\Pi = 0)$, where the subscript signifies random realizations of $S$ due to permutations $\Pi$. The basic approach here is to (i) choose $\alpha = \min_i p_i$, (ii) compute the first two moments of $S_\Pi$ under the bivariate normal assumption for pairs of $z$ statistics, (iii) approximate $P(S_\Pi = 0)$ using these two moments and an appropriate distributional approximation for $S_\Pi$. The final result is approximate FWER control, but computed much faster than using actual permutation.

For the moments, we note that for $m$ statistics, $E(S_\Pi) = \sum_{i=1}^m E(I_{i,\Pi}) \approx m\alpha$. For the second moment, we have $\text{var}(S_\Pi) = \sum_i \sum_{i'} \text{cov}(I_{i,/}, I_{i',\Pi})$, where $\text{cov}(I_{i',\Pi}, I_{i',\Pi}) = P(I_{i,\Pi} = 1, I_{i',\Pi} = 1) - \alpha^2$. For threshold $z_\alpha = \Phi^{-1}(\alpha/2)$, Figure 5 illustrates the joint rejection region $\{I_i = 1, I_{i'} = 1\}$ for a pair of $z$ statistics with correlation $\rho = 0.6$. These probabilities must be computed for the $\binom{m}{2}$ distinct pairs $\{i \neq i'\}$, with $\rho_{i,i'} = r(x_i, x_{i'})$. For large $m$, numerical integration of the bivariate normal density for the rejection region becomes computationally intensive, even using $R$ functions such as *pbinorm*() of the $VGAM$ package that have been optimized for this purpose. Here we gain computational efficiency using the approximation from Cox and Wermuth (1991). For threshold $z_\alpha$,

$$P(Z_1 > z_\alpha, Z_2 > z_\alpha) \approx$$

$$\Phi(-z_\alpha)\{\Phi(\xi(z_\alpha, \rho)) - \frac{\rho^2}{2(1-\rho^2)}\xi(z_\alpha, \rho)\phi(\xi(z_\alpha, \rho))\sigma^2\}, \quad (5)$$

where $\xi(z_\alpha, \rho) = \frac{\rho\mu - z_\alpha}{\sqrt{1-\rho^2}}$, $\mu = \phi(z_\alpha)/\Phi(-z_\alpha)$, and $\sigma^2 = 1 + z_\alpha\mu - \mu^2$ (the last equation correcting a typo in Cox and Wermuth (1991). From (4) and trivial symmetries we obtain the final probability, expressed visually as the four regions shown in Figure 5 for an illustrative pair of $z$ statistics.

Using the approximations above proceeds as noted by plugging in the correlations of each pair of markers to substitute as correlations for $z$ and is reasonably accurate, but slightly underestimates $\mathrm{var}(S_\Pi)$. To see why, note that even if $r(x_1, x_2) = 0$, $r_{1,\Pi}$ and $r_{2,\Pi}$ are not independent, as the fact that $x_1$ and $x_2$ are orthogonal linear predictors of $y$ implies that $r^2_{1,\Pi} + r^2_{2,\Pi} \leqslant 1$. The consequence of the constraint can be easily estimated for the situation with $r(x_1, x_2) = 0$ using the standard beta distributional approximation (Zhou et al., 2013) with $r^2_{1,\Pi} \dot{\sim} \mathrm{Beta}(\frac{1}{2}, \frac{n-2}{2})$ and $r^2_{2,\Pi} | r^2_{1,\Pi} \dot{\sim} (1 - r^2_{1,\Pi})\mathrm{Beta}(\frac{1}{2}, \frac{n-3}{2})$. If $B_n(a) = P(r^2_{1,\Pi} > a)$ and $F_n(a) = P(r^2_{1,\Pi} > a, r^2_{2,\Pi} > a)$ based on the beta distributions as described, then we compute the "deficit" in joint probability of exceedance compared to independence as $F_n(a) - B^2_n(a)$. This deficit is then used as an offset for each covariance of indicators in estimating $\mathrm{var}(\mathrm{S}_\Pi)$.

Finally, using the estimates $\hat{E}(S_\Pi)$, $\widehat{\mathrm{var}}(S_\Pi)$, we model the distribution of $S_\Pi$. If the variance exceeds the mean (overdispersion), a simple and effective approach is to model $S_\Pi$ as a mixture of two distributions: i) approximately Poisson with probablity $\pi_1$ and ii) a point mass assuming the value $m(1 - \pi_1)$ with probability $\alpha$. Here $\pi_1$ is chosen so that the mixture matches the overall mean and variance. Finally the $p$-value is estimated as $P(S_\Pi > 0) \approx 1 - e^{-m\pi_1\alpha}(1 - \alpha)$.

For a dataset consisting of a single marker and multiple traits, we can use the methods in the previous subsections to instead compute the FWER-controlling approximate $p$-value across the $t$ traits simply by substituting the $t \times t$ trait correlation matrix in place of the genotype correlation matrix.

*The correlation between $r_\Pi$ at two markers for two traits, under permutation*

**Result 2**

Using $r_{i,k,\Pi}$ to denote the random correlation between the $i$th marker and the $k$th trait,

$$\operatorname{corr}(r_{i,k,\Pi}, r_{i',k',\Pi}) = r(x_i, x_{i'})r(y_k, y_{k'}). \tag{6}$$

*Proof.* First we note that

$$\sum_j \sum_{j'} y_{k,j} y_{k',j'} = (\sum_j y_{j,k})(\sum_{j'} y_{k',j'}) = 0. \tag{7}$$

The result in (6) also holds for any permutation of the indices, so taking expectations on both sides for a random $\Pi$ gives

$$\begin{aligned}\sum_j \sum_{j'} E(y_{k,\Pi[j]} y_{k',\Pi[j']}) = 0 &= \sum_j E(y_{k,\Pi[j]} y_{k',\Pi[j]}) + \sum_{j \neq j'} E(y_{k,\Pi[j]} y_{k',\Pi[j']}) \\ &= r(y_k, y_{k'}) + (n^2 - n)E(y_{k,\Pi[j]} y_{k',\Pi[j']}),\end{aligned} \tag{8}$$

implying $E_{j \neq j'}(y_{k,\Pi[j]} y_{k',\Pi[j']}) = -r(y_k, y_{k'})/(n^2 - n)$, and where the final expectation holds for any $j \neq j'$. Plugging these results into the two-trait analogue of (3) gives

$$\begin{aligned}\operatorname{cov}(r_{i,k,\Pi}, r_{i',k'\Pi}) &= \sum_j x_{1j} x_{2j} E(y_{k,\Pi[j]} y_{k',\Pi[j]}) + \sum_{j \neq j'} x_{1j} x_{2j'} E(y_{k,\Pi[j]} y_{k'\Pi[j']}) \\ = \sum_j x_{1j} x_{2j} (\frac{r(y_k, y_{k'})}{n}) &+ \sum_{j \neq j'} x_{1j} x_{2j'} (\frac{-r(y_k, y_{k'})}{n(n-1)}) = \frac{r(x_i, x_{i'}) r(y_k, y_{k'})}{n-1}\end{aligned} \tag{8}$$

where the last equality follows by a comparison with eq. (4), and we divide the last term by $1/(n-1)$ to obtain the final correlation.

The remarkably simple result occurs due to the scheme of permuting entire columns of $Y$, which is desirable to preserve trait-trait correlation structure.

*Correcting family-wise error across multiple markers and traits*

Following Result 2, it is straightforward to adapt the previous methods to the situation in which we have multiple traits arranged in a $t \times n$ matrix $Y$.

When computing the family-wise error across *all* traits and markers, we expand the definition $S = \sum_i \sum_k I_{ik}$ where $I_{ik} = I[p_{ik} \leqslant \alpha]$, and $p_{ik}$ is the $p$-value for the $i$th marker and $k$th trait. We apply the same reasoning as in subsection 2.4 above, but need to sum over the $m^2 \times t^2$ elements in the covariance matrix. Using $\rho_X$ to denote the $m \times m$ correlation matrix of the rows of $X$, and $\rho_Y$ to denote the $t \times t$ correlation matrix among the rows of $Y$, we finally compute the $(mt) \times (mt)$ Kronecker product $\rho_{XY} = \rho_X \otimes \rho_Y$, and use $\rho_{XY}$ directly in applying the methods of subsection 2.3 to FWER correction for the $mt$ $p$-values.

We refer to this $p$-value after correcting for all marker-trait comparisons as the *total* FWE-controlling $p$-value. It measures the evidence that any marker is associated with any trait.

*A speedup of the exceedance variance calculation*

The Cox-Wermuth bivariate normal approximation offers a substantial speedup in comparison to full numeric integration, but can be intensive when $m$ and $t$ are large. The variance of $S_\Pi$ depends only on the overall sum of the associated covariance matrix, so it is sufficient to work with a weighted grid of original data covariances (e.g. for $X$) and convert these grid values to covariances of indicators $I$. Specifically, the histogram of $m^2$ $X$ correlations is computed across a grid of bin midpoints (maximum default 50 values), with weights according to the number of correlations falling in the midpoint of each bin. Then only these 50 values need to be carried forward in calculaton of the covariance of indicators as shown in subsection .

*Quadratic form association evidence across traits and markers*

The previous focus on FWER control is sensible in situations where a single marker is expected to provided dominant evidence of association. However, studies have shown evidence of mutiple causal markers per gene (e.g. in eQTL studies, Jansen et al. (2017)), for which aggregation across multiple markers may improve detection power. The quadratic form approach Zhou et al. (2013) is used to provide summarized evidence across multiple markers as the sum of squared score statistics, which after scaling steps is equivalent to summing squared marker-trait correlations across multiple markers.

For appropriately standardized $X$ (row-centered and scaled) and single $1 \times n$ trait $y$, we use the statistic $Q = yAy^T$, where $A = X^T X$. The methods Zhou et al. (2013) are used to provide the first four moments of $Q_\Pi$, for which $p$-values are obtained with a Pearson family density approximation having the same moments as $Q_\Pi$. Similarly, for a single marker $x_i$, the same approach can be used for summarized evidence across the traits as $x_i Y^T Y x_i^T$.

For an overall aggregate summary across all traits and markers, a natural statistic is the overall sum of squared score statistics

$$Q_{total} = \sum_k Q_k = \sum_k y_k A y_k^T = \sum_{k=1}^{t} \sum_{i=1}^{n} \sum_{j=1}^{n} a_{ij} y_{ki} y_{kj} = \sum_{i=1}^{n} \sum_{j=1}^{n} a_{ij} b_{ij},$$

where $b_{ij} = \sum_k y_{ki} y_{kj}$. It is not immediately clear how to obtain permutation moments for $Q_{total}$, and the $Q_k$ terms are in general correlated, as the permutation keeps the columns of trait values intact. However, a device similar to that used in Zhou et al. (2013) enables conversion of $Q_{total}$ into a form amenable to using the Semiatycki moments developed for statistics designed to detect space-time clustering Siemiatycki (1978). Note that we have scaled $X$ so that each row and column of $A = X^T X$ sums to zero. We define the $m \times m$ matrix $C = A - diag(A)$, and the $m \times m$ matrix D with elements $d_{ij} = -\frac{1}{2} \sum_k (y_{ki} - y_{kj})^2$. Define $W = \sum_i \sum_j c_{ij} d_{ij}$. $C$ and $D$ have the necessary characteristics for the Semiatycki moment calculation: (i) they are symmetric, and (ii) they have zero diagonals. It is easy to verify that column permutation of the $Y$ matrix is equivalent to simultaneous permutation of the rows and columns of $D$. Thus the permutation moments of $W$ may be computed using the methods Zhou et al. (2013). It remains to show that $W = Q_{total}$, and in fact they remain equal under each permutation, so that $W_\Pi = Q_{total,\Pi}$. We have $\sum_i \sum_j c_{ij} d_{ij} = \sum_i \sum_j a_{ij} \{-\frac{1}{2} \sum_k (y_{ki} - y_{kj})^2\}$ because each $d_{ii} = 0$, and expanding the terms in braces gives

$$\sum_i \sum_j c_{ij} d_{ij} = \underbrace{\sum_i \sum_j a_{ij} \sum_k y_{ki} y_{kj}}_{Q_{total}} - \frac{1}{2} \sum_i \sum_j a_{ij} \sum_k y_{ki}^2 - \frac{1}{2} \sum_i \sum_j a_{ij} \sum_k y_{kj}^2.$$

Each of the last two terms is zero, because for instance $\sum_i \sum_j a_{ij} \sum_k y_{ki}^2 = \sum_i \sum_k (y_{ki}^2)(\sum_j a_{ij})$ and $\sum_j a_{ij} = 0$.

We refer to the $p$-value based on $Q_{total}$ as the *total* quadratic form $p$-value. It measures the aggregate evidence that the markers are associated with the traits.

*Eigenvalue-conditional moments for the quadratic form*

*Motivation*

The four moments shown in subsection  for row totals, column totals, and the overall total are *exact* under permutation of columns of $Y$ (say) in relation to $X$, but their computation is $O(n^3)$ even under the fastest method available Zhou et al. (2013). For moderate to large $n$, another approach is to extend the reasoning for moments of summed squared correlations Zhou et al. (2013), which was also shown to be a quadratic form. For an arbitrary vector $y$ (a column from $Y$) we use the relation $y^T X^T X y = y^T P^T \Lambda P y$ using an eigendecomposition $X^T X = P^T \Lambda P$. Then $\sum_i corr(X_{i.}, y)^2 = \sum_i r_{ij}^2 = \sum_i \lambda_i corr(P_{.i}, y)^2 = \sum_i \lambda_i u_i^2$, where $P_{.i}$ is the $i$th column eigenvector of $X^T X$. In contrast to correlations among the $r^2$ values, the vectors of $P$ are orthogonal and the values $corr(P_., y)^2$ amenable to moment approximation under permutation. It is well known that squared correlations $corr(x, y)^2$ follow a beta density when either $x$ or $y$ is normal. As discussed in Zhou et al. (2013), the orthogonality of columns of $P$ creates slight dependence of correlations, and here we will use the observed eigenvalues of $X^T X$ for our calculations. Thus our moments derived below may be viewed as *eigenvalue-conditional* for Gaussian data.

In practice, we use these eigenvalue conditional moments to approximate the permutation distributions of the summed statistics. The rationale is that permutation of course preserves the eigenvalues of $X$ and $Y$, and beta densities are a good approximation to the permutation distribution of squared correlations Zhou et al. (2013).

*Derivation*

Let $S = \sum_i \lambda_i u_i^2$. For a single squared Pearson correlation $A = u_i^2$, we use the null approximation $A \sim Beta(1/2, (n-2)/2)$. Some moments and central moments of $A$ follow from the fact that the $k$th raw moment is $\Pi_{l=0}^{k-1} \frac{1/2+l}{(n-1)/2+l}$: $E(A) = 1/(n-1)$ (which is also an exact result under permutation), $E(A^2) = \frac{3}{(n-1)(n+1)}$, $\text{var}(A) = \frac{2(n-2)}{(n-1)(n^2-1)}$, $E(A^3) = \frac{15}{(n+3)(n^2-1)}$, $skewness(A) = \frac{4(n-3)\sqrt{(n+1)/2}}{(n+3)\sqrt{n-2}}$, and $E(A^4) = \frac{105}{(n-1)(n+1)(n+3)(n+5)}$. Due to orthogonality of columns of $P$, the $\{u_i^2\}$ are somewhat negatively correlated, and the impact of the correlations may be non-trivial if $m$ is large and the eigenvalues are not dominated by a few values. We have $corr(u_i^2, u_j^2) = \frac{-1}{n-2}$, which can be obtained using the technique of successive correlated beta random variables Zhou et al. (2013) $u_i^2 = A \sim Beta(1/2, (n-2)/2)$, and $u_j^2 = B(1-A)$, where $B$ is independent of $A$ and $B \sim Beta(1/2, (n-3)/2)$. This approach can be used for successive beta random variables to represent the third (and fourth) in a succession of squared correlations. From this approach/results and the fact that $\sum_i \lambda_i = m$ we obtain $E(S) = E(\sum_i \lambda_i u_i^2) = \frac{m}{n-1}$ and after some cancellation $\text{var}(S) = \sum_i \sum_j \text{cov}(u_i^2, u_j^2) = (\sum_i \lambda_i^2 - \frac{m^2}{n-1})(\frac{2}{n^2-1})$.

To obtain the third moment of $S$, we extend the correlated beta technique. We have

$$E(S^3) = \sum_i \sum_j \sum_k \lambda_i \lambda_j \lambda_k E(u_i^2 u_j^2 u_k^2).$$

For the inner terms, if $i = k = j$, then we use the third moment from above. If $i = j, j \neq k$, we use the $A, B$ notation from above. We have $E(u_i^4 u_j^2) = E(A^2 B(1-A)) = E(B)(E(A^2) - E(A^3)) = \frac{1}{n-2}(\frac{3}{n^2-1} - \frac{15}{(n+3)(n^2-1)}) = \frac{3}{(n+3)(n^2-1)}$. If $i \neq j, j \neq k, i \neq k$, then $E(u_i^2 u_j^2 u_k^2) = E(AB(1-A)C(1-A-B(1-A))) = E(C)E(A(1-A)^2)E(B(1-B))$, and $E(B) = \frac{1}{n-2}, E(C) = \frac{1}{n-3}$, etc. After some steps, we have $E(A(1-A)^2) = \frac{n(n-2)}{(n-1)(n+1)(n+3)}$ and $E(B - B^2) = \frac{n-3}{n(n-2)}$. Substituting all these values produces cancellations and $\underbrace{E(u_i^2 u_j^2 u_k^2)}_{i \neq j, j \neq k, i \neq k} = \frac{1}{(n+3)(n^2-1)}$. Finally, we substitute the various terms in summations and note additional cancellations, giving $E(S^3) = \frac{1}{(n^2-1)(n+3)}(m^3 + 6m \sum_i \lambda_i^2 + 8 \sum_i \lambda_i^3)$, from which the skewness of $S$ is easily calculated.

The fourth moment requires careful attention to keep track of terms. We write

$$E(S^4) = \sum_i \sum_j \sum_k \sum_l \lambda_i \lambda_j \lambda_k \lambda_l E(u_i^2 u_j^2 u_k^2 u_l^2)$$

and for notational convenience for below we suppose that the subscripts $i$, $j$, $k$, $l$ are all distinct unless otherwise stated. Then

$$\begin{aligned}
\mathrm{E(S^4)} &= \textstyle\sum_i \lambda_i^4 E(u_i^8) + \underbrace{\sum_i \sum_j}_{i \neq j} \lambda_i \lambda_j^3 E(u_i^2 u_j^6) + \underbrace{\sum_i \sum_j}_{i \neq j} \lambda_i^2 \lambda_j^2 E(u_i^4 u_j^4) \\
&+ \underbrace{\sum_i \sum_j \sum_k}_{i \neq j, j \neq k, i \neq k} \lambda_i \lambda_j \lambda_k^2 E(u_i^2 u_j^2 u_k^4) + \underbrace{\sum_i \sum_j \sum_k \sum_l}_{i \neq j, i \neq k, i \neq l, j \neq k, j \neq l, k \neq l} \lambda_i \lambda_j \lambda_k \lambda_l E(u_i^2 u_j^2 u_k^2 u_l^2) \\
&= E(u_i^8) \underbrace{\sum_i \lambda_i^4}_{①} + \underbrace{E(u_i^2 u_j^6)}_{i \neq j} \underbrace{\underbrace{\sum_i \sum_j}_{i \neq j} \lambda_i \lambda_j^3}_{②} + \underbrace{E(u_i^4 u_j^4)}_{i \neq j} \underbrace{\underbrace{\sum_i \sum_j}_{i \neq j} \lambda_i^2 \lambda_j^2}_{③} \\
&+ \underbrace{E(u_i^2 u_j^2 u_k^4)}_{i \neq j, j \neq k, i \neq k} \underbrace{\underbrace{\sum_i \sum_j \sum_k}_{i \neq j, j \neq k, i \neq k} \lambda_i \lambda_j \lambda_k^2}_{④} + \underbrace{E(u_i^2 u_j^2 u_k^2 u_l^2)}_{i \neq j, i \neq k, i \neq l, j \neq k, j \neq l, k \neq l} \underbrace{\underbrace{\sum_i \sum_j \sum_k \sum_l}_{i \neq j, i \neq k, i \neq l, j \neq k, j \neq l, k \neq l} \lambda_i \lambda_j \lambda_k \lambda_l}_{⑤}
\end{aligned}$$

We first handle the expectations, which do not depend on the specific indexes $i$, $j$, $k$, $l$. $E(u_i^8) = E(A^4) = \frac{105}{(n-1)(n+1)(n+3)(n+5)}$. $\underbrace{E(u_i^2 u_j^6)}_{i \neq j} = E(A^3 B(1-A)) = E(B) E(A^3(1-A)) = E(B)(E(A^3) - E(A^4)) = \frac{15}{(n-1)(n+1)(n+3)(n+5)}$. $\underbrace{E(u_i^4 u_j^4)}_{i \neq j} = E(A^2(B(1-A))^2) = E(A^2(1-A)^2 B^2) = E(B^2) E(A^2(1-A)^2) = \frac{1}{n-2} \frac{3}{n} (E(A^2) - 2E(A^3) + E(A^4))$. Plugging in the various expectations leads to numerous cancellations, and finally $\underbrace{E(u_i^4 u_j^4)}_{i \neq j} = \frac{9}{(n-1)(n+1)(n+3)(n+5)}$.

$\underbrace{E(u_i^2 u_j^2 u_k^2 u_l^2)}_{i \neq j, i \neq k, i \neq l, j \neq k, j \neq l, k \neq l} = E(A^2 B(1-A) C(1 - B(1-A) - A)) = E(C(A^2(1-A)^2 B(1-B))$. We have $E(C) = \frac{1}{n-3}$ and $E(B(1-B)) = \frac{n-3}{n(n-2)}$, and $E(A^2(1-A)^2) = E(A^2 - 2E(A^3) + E(A^4) = ... = \frac{3n(n-2)}{(n-1)(n+1)(n+3)(n+5)}$. Plugging in all terms gives $\underbrace{E(u_i^2 u_j^2 u_k^2 u_l^2)}_{i \neq j, i \neq k, i \neq l, j \neq k, j \neq l, k \neq l} = \frac{3}{(n-1)(n+1)(n+3)(n+5)}$.
The last expectation is $\underbrace{E(u_i^2 u_j^2 u_k^2 u_l^2)}_{i \neq j, i \neq k, i \neq l, j \neq k, j \neq l, k \neq l} = E(AB(1-A)(C(1-B(1-A)-A))D(1-C(1-B(1-A)-A) - B(1-A) - A))$, which simplifies to $E(A(1-A)^3 B(1-B)^2 C(1-C) D) = E(D) E(C(1-C)) E(B(1-B)^2) E(A(1-A)^3)$. We have $E(D) = \frac{1}{n-4}$, $E(C(1-C)) = \frac{1}{n-3} \frac{n-4}{n-1}$. $E(B(1-B)^2) = ... = \frac{1}{n-2} \frac{(n-3)(n-1)}{n(n+2)}$. $E(A(1-A)^3) = E(A - 3A^2 + 3A^3 - A^4)$, and

collecting terms with common denominator gives after cancellations $\underbrace{E(u_i^2 u_j^2 u_k^2 u_l^2)}_{i\neq j, i\neq k, i\neq l, j\neq k, j\neq l, k\neq l} = \frac{1}{(n-1)(n+1)(n+3)(n+5)}$.

The next task is to calculate the summations. We have

$$①=\sum_i \lambda_i^4,$$

$$②=\underbrace{\sum_i \sum_j}_{i\neq j} \lambda_i \lambda_j^3 = 4\Big(\sum_i \sum_j \lambda_i \lambda_j^3 - ①\Big) = 4\Big(m\sum_i \lambda_i^3 - ①\Big),$$

$$③=\underbrace{\sum_i \sum_j}_{i\neq j} \lambda_i^2 \lambda_j^2 = 3\Big(\sum_i \sum_j \lambda_i^2 \lambda_j^2 - ①\Big) = 3\Big((\sum_i \lambda_i^2)^2 - ①\Big),$$

$$④=\underbrace{\sum_i \sum_j \sum_k}_{i\neq j, j\neq k, i\neq k} \lambda_i \lambda_j \lambda_k^2 = 6\Big(\sum_i \sum_j \sum_k \lambda_i \lambda_j \lambda_k^2 - ③/3 - ②/2 - ①\Big) = 6\Big(m^2 \sum_i \lambda_i^2 - ③/3 - ②/2 - ①\Big),$$

$$⑤ = m^4 - (① + ② + ③ + ④),$$

where the last value reflects the fact that $\sum_i \sum_j \sum_k \sum_l \lambda_i \lambda_j \lambda_k \lambda_l = m^4$. The coefficient terms are multinomial coefficients representing the number of ways in which the indexes can be chosen. After substituting the summations and the expectations from earlier, the final fourth raw moment is

$$E(S^4) = \frac{1}{(n-1)(n+1)(n+3)(n+5)}\Big(m^4 + 48\sum_i \lambda_i^4 + 32m\sum_i \lambda_i^3 + 12(\sum_i \lambda_i^2)^2 + 12m^2\sum_i \lambda_i^2\Big)$$

All of the above has been developed for the summation across $m$ markers. The identical formulas apply for summation across traits, substituting the eigenvalues of $Y^TY$ and $t$ for $m$, etc. Estimating the permutation moments for the total sum $Q_{total}$ requires an additional approximation. First, we note that $Q_{total}$ can be, in a manner similar to above, similarly decomposed as a sum of $Q_{total} = \sum_i \sum_k \lambda_i \eta_i corr(P_{.i}, Q_{.k})^2$, where $\lambda_i$ is the $i$ eigenvalue of $X^TX$ and $\eta_k$ is the $k$th eigenvalue of $Y^TY$, and $P$ and $Q$ are the respective matrices of eigenvectors.

#### References

Cox, D. R. and Wermuth, N. (1991). A simple approximation for bivariate and trivariate normal integrals. *International Statistical Review/Revue Internationale de Statistique* pages 263–269.

Jansen, R., Hottenga, J.-J., Nivard, M. G., Abdellaoui, A., Laport, B., de Geus, E. J., Wright, F. A., Penninx, B. W., and Boomsma, D. I. (2017). Conditional eqtl analysis reveals allelic heterogeneity of gene expression. *Human molecular genetics* **26,** 1444–1451.

Siemiatycki, J. (1978). Mantel's space-time clustering statistic: computing higher moments and a comparison of various data transforms. *Journal of statistical Computation and Simulation* **7,** 13–31.

Zhou, Y.-H., Barry, W. T., and Wright, F. A. (2013). Empirical pathway analysis, without permutation. *Biostatistics* **14,** 573–585.

Zhou, Y.-H., Mayhew, G., Sun, Z., Xu, X., Zou, F., and Wright, F. A. (2013). Space–time clustering and the permutation moments of quadratic forms. *Stat* **2,** 292–302.
